# Identifying Representative Cell Line Models for TNBC Chemotherapy Drug Resistance via Systems Biology and Bioinformatics

**DOI:** 10.1101/2024.05.22.595413

**Authors:** Shuai Shao, Lang Li

## Abstract

Cancer cell lines, derived from tumors, have become essential tools in life science research and are commonly employed as experimental model systems in cancer research. However, researchers often overlook the similarities between clinical samples when selecting cell line models, potentially impacting the validity and applicability of their findings. In the context of triple-negative breast cancer (TNBC) chemotherapy drug resistance, our study aims to provide guidance for selecting appropriate cell line models by employing a combination of systems biology and bioinformatic approaches. These approaches, including hierarchical clustering analysis, Spearman’s rank correlation, and single sample gene set enrichment analysis (ssGSEA), allowed us to identify the most representative cell models that correspond to poor chemotherapy responders among TNBC patients.

## Introduction

Breast cancer is the most prevalent cancer globally, accounting for 12.5% of all new annual cancer cases. In 2022, the American Cancer Society estimated that approximately 13% of U.S. women (1 in 8) would develop invasive breast cancer during their lifetime. They projected 287,850 new cases of invasive breast cancer and 51,400 new cases of non-invasive (in situ) breast cancer to be diagnosed in women in the U.S. Triple-negative breast cancer (TNBC) is a highly aggressive subtype of breast cancer that disproportionately impacts younger women, particularly those of African descent [1]. TNBC makes up 10-20% of all breast cancer cases and is characterized by the absence of estrogen and progesterone receptors, as well as the lack of human epidermal growth factor receptor 2 overexpression [2].

Chemotherapy remains a primary therapeutic strategy for managing TNBC, which is known for its biological aggressiveness. Chemo therapeutic strategies for TNBC management involve targeting DNA repair complexes (e.g., platinum compounds), microtubules (e.g., taxanes), and cell proliferation (e.g., anthracycline-containing regimens)[3]. Numerous neoadjuvant studies have investigated the potential advantages of combining innovative chemotherapeutic agents with standard chemotherapy, encompassing anthracyclines, taxanes, antimetabolites, platinum-based compounds, and new microtubule stabilizing agents[2]. Third-generation chemotherapy strategies employing dose-dense or metronomic polychemotherapy, akin to those provided to other high-risk patients, rank among the most potent approaches currently accessible for treating both early-stage and advanced disease[4] .

In recent years, significant progress in understanding the molecular characteristics of TNBC has led to the development of potential new therapeutic options for TNBC patients. Some of these novel therapies include immunotherapy, such as atezolizumab (Tecentriq) [5], which has shown promising results in combination with nab-paclitaxel in advanced triple-negative breast cancer. Additionally, PARP inhibitors like olaparib [6], niraparib (Zejula), and talazoparib (Talzenna) [7] have emerged as valuable options, particularly when combined with chemotherapy[8] agents such as paclitaxel for first- or second-line treatment in metastatic triple-negative breast cancer patients. Another innovative approach is the use of antibody-drug conjugates (ADCs) [9, 10], such as sacituzumab govitecan (Trodelvy), which selectively deliver cytotoxic agents to cancer cells while minimizing damage to healthy cells. Furthermore, targeted therapies based on specific molecular mechanisms [11], like PI3K inhibitors (e.g., alpelisib, buparlisib) and mTOR inhibitors (e.g., everolimus), have gained traction, offering new avenues for TNBC treatment. These advances highlight the potential for novel, targeted, and personalized therapies that build on the growing understanding of TNBC’s molecular landscape.

The first human cancer cell line, known as HeLa, was discovered and established over sixty years ago [12]. Since its inception, it has become widely popular among research institutions globally. Cancer cell lines, derived from tumors, have become essential tools in life science research, owing to the advantages of cell culture. Consequently, cancer cell lines are commonly employed as experimental model systems in cancer research.

However, researchers often overlook the similarities between clinical samples when selecting cell line models for their experiments. For instance, PubMed citations reveal that 40.2% of research in metastatic breast cancer utilized the MDA-MB-231 cell line model. Nevertheless, Bin Chen et al.2 discovered that MDA-MB-231 exhibited lower similarity to patient samples of basal-like metastatic breast cancer compared to other cell lines [13]. In another example, Domcke et al. [14] investigated the genomic profiles of 47 ovarian cancer cell lines and ovarian cancer tumor samples. They found that several underutilized cell lines more closely resembled high-grade serous ovarian tumor samples than the more popular cell lines. This highlights the need for researchers to carefully consider the relevance and similarity of cell line models to clinical samples when conducting cancer research, as this could significantly impact the validity and applicability of their findings.

In vitro human cell line models are extensively employed in studies investigating chemotherapy drug resistance in triple-negative breast cancer (TNBC). Numerous gene drivers and pathways have been identified as contributing factors to TNBC chemotherapy drug resistance [15-18]. However, researchers often overlook the importance of matching the genomic profiles of patients and cell lines. Utilizing cell line models that do not accurately represent clinical situations can lead to unreliable predictions of clinical responses. Our work aims to provide guidance for selecting appropriate cell line models in the context of TNBC chemotherapy drug resistance. A better cell line model facilitates a more efficient exploration of the mechanisms underlying chemotherapy resistance in TNBC patients. Understanding these mechanisms can lead to the identification of improved drug treatment targets. Furthermore, the methods implemented in our studies offer insights into using cell line models to determine similarities among patients in other research areas.

In our study, we employed a combination of systems biology and bioinformatic approaches, including hierarchical clustering analysis, Spearman’s rank correlation, and gene set enrichment analysis (GSEA) based on gene expression levels and pathway activation patterns. These methods allowed us to identify suitable cell models that correspond to poor chemotherapy responders among TNBC patients. This comprehensive approach will enhance the selection of appropriate cell line models and contribute to a more accurate understanding of TNBC chemotherapy drug resistance, ultimately leading to better-targeted therapies for patients.

## Result

### Hierarchical clustering analysis by genomic profiles comparation

Our study aims to identify the cell lines that best represent pre-treatment chemo drug poor responders and post-chemo poor responders, respectively. To achieve this, firstly we grouped patients into the resistant and drug-sensitive groups at baseline and post-chemo time points separately and performed differential gene expression analysis. We identified significantly upregulated or downregulated genes in resistant vs. sensitive patients using the screening criteria for differentially expressed genes (fold change greater than 1.5 or less than -1.5, adjusted p-value less than 0.01, Table 6 and Table 7). We then used these gene expression data to perform hierarchical clustering analysis on patients and TNBC cell lines from the CCLE. Based on hierarchical clustering analysis, subgroups of samples with similar gene expression patterns can be identified. Heatmaps displaying the clustering results for baseline samples are shown in Figure 1A, while Figure 2A presents the post-chemotherapy samples. In both clustering outcomes, clear patterns and distinctions are evident between good responders and poor responders. Furthermore, some TNBC cell lines are grouped in the same subgroup as the TNBC poor responders, demonstrating their potential as representative models for these patients.

**Figure 1.**
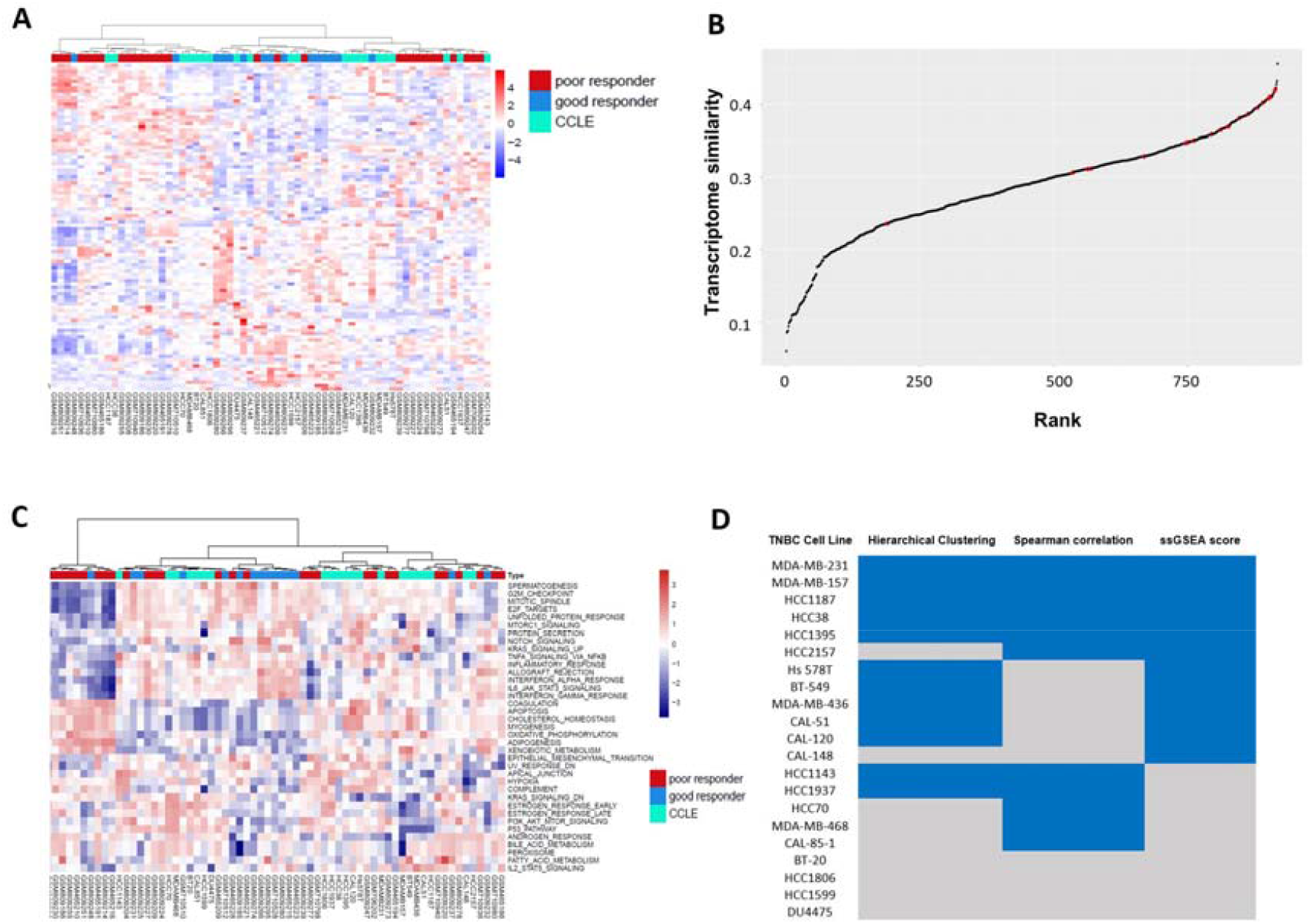
Genomic similarity analysis between TNBC patients (baseline) and CCLE cell line model. (A) Hierarchical clustering analysis. (B) 916 CCLE cell lines are ranked according to Spearman’s rank correlation value with TNBC poor responder samples. (C) Heatmap of ssGSEA scores for the 50 MSigDB hallmark gene sets across TNBC patient samples and TNBC cell lines. (D) Summary table of the most representative cell line models from these three analysis methods. Cell lines marked in blue indicate significant representativeness for TNBC poor responders.

**Figure 2.**
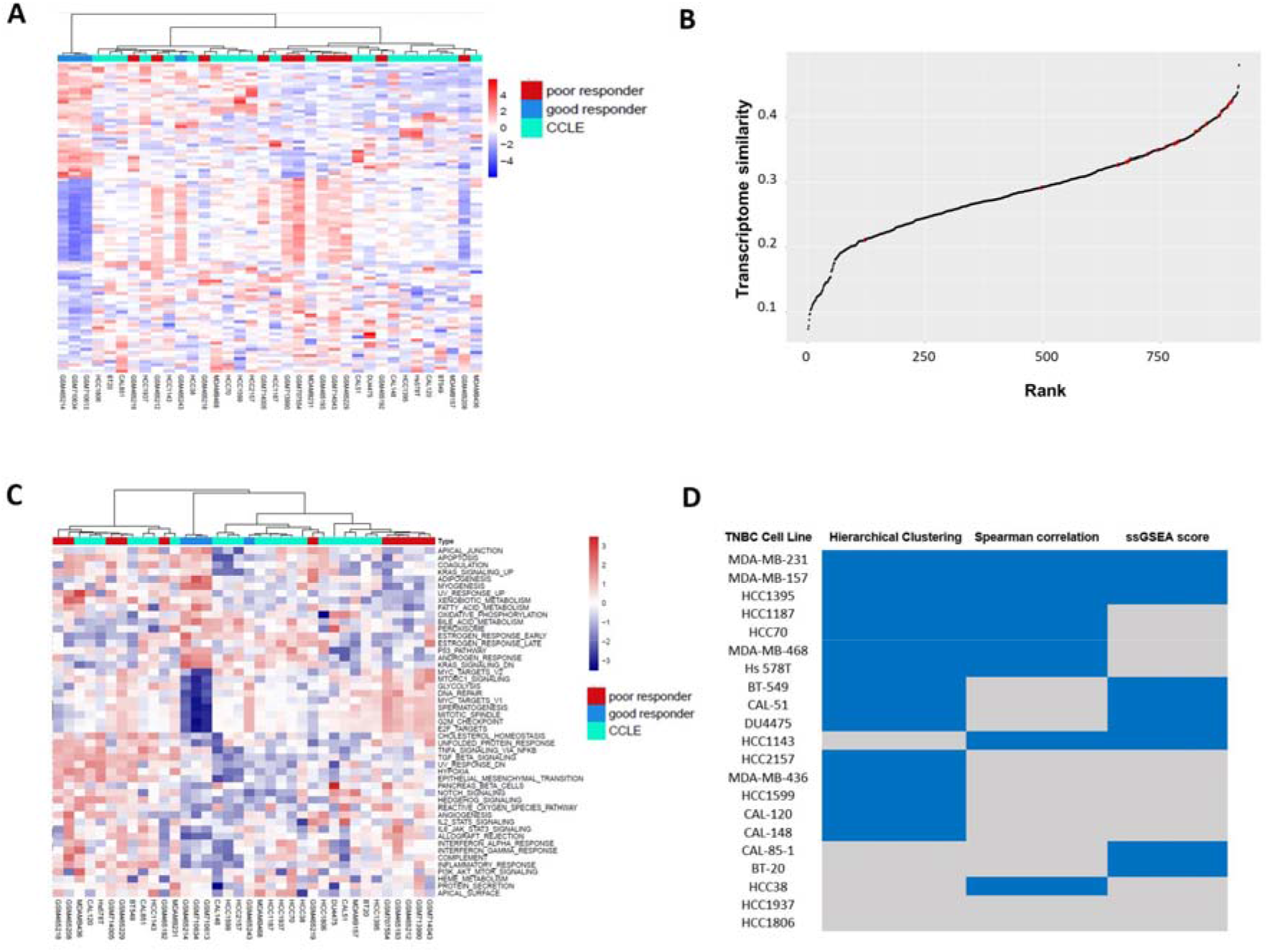
Genomic similarity analysis between TNBC patients (Post chemotherapy) and CCLE cell line model. (A) Hierarchical clustering analysis. (B) 916 CCLE cell lines are ranked according to Spearman’s rank correlation value with TNBC poor responder samples. (C) Heatmap of ssGSEA scores for the 50 MSigDB hallmark gene sets across TNBC patient samples and TNBC cell lines. (D) Summary table of the most representative cell line models from these three analysis methods. Cell lines marked in blue indicate significant representativeness for TNBC poor responders.

### Correlating cell lines with TNBC poor responder

Spearman correlation analysis was performed between the 916 CCLE cell lines and TNBC poor responders using the 2,000 most varied genes. Cell lines were ranked based on their average correlation values. In the correlation plots (Figure 1B for correlation with baseline samples and Figure 2B for correlation with post-chemotherapy samples), TNBC samples are labeled in red, while cell lines for other cancer types are labeled in black. with those having the highest values being the most representative of the patient population. HCC70 and DU4475 are the two TNBC cell lines with the highest and lowest transcriptome similarity, respectively (Spearman rank correlations are 0.42, rank 5 and 0.24, rank 729 in Table 8), with baseline TNBC poor responders. HCC1143 and DU4475 are the two TNBC cell lines with the highest and lowest transcriptome similarity, respectively (Spearman rank correlations are 0.42, rank 17 and 0.21, rank 796 in Table 9), with post-chemo TNBC poor responders.

### Pathway similarity evaluation by ssGSEA score

We utilized ssGSEA scores of the 50 MSigDB hallmark gene sets to assess the similarity between TNBC cell lines and TNBC poor responder samples at the pathway activation level. We generated heatmaps using hierarchical clustering analysis (Figure 1C for baseline samples and Figure 2C for post-chemo samples) based on the ssGSEA scores to visualize the similarity between cell lines and TNBC poor responder patient samples. The results of the hierarchical clustering heatmaps revealed distinct clusters of TNBC cell lines and TNBC patient poor responder samples with similar MSigDB hallmark gene pathway activation profiles. Furthermore, by examining these clusters, we could identify cell lines that are more similar to TNBC-poor responders in terms of pathway activation level (Figures 1D and 2D).

## Discussion

The selection of accurate and representative cell line models is crucial for the success of medical translation. An accurate cell line model can closely mimic the molecular and cellular characteristics of the target patient population, enabling researchers to better understand disease mechanisms, identify potential drug targets, and evaluate the efficacy and safety of new therapeutic interventions. Our study analyzes public genomic profiles to assess the similarity between breast cancer cell lines and TNBC patient samples. We aim to find the most representative cell lines for TNBC poor responders at both baseline and post-chemo time points. Our evaluation considers not only the similarity at the gene expression level but also the pathway similarity. We applied hierarchical clustering and Spearman correlation analysis to the gene expression data. Distinct patterns and differences are discernible between good responders and poor responders. Approximately ten cell lines cluster in each study with the poor responder group (Figure 1D and Figure 2D).

Transcriptome correlation analysis (TC analysis) is proven to be an effective approach to evaluate cell lines for research purpose [19]. Higher average correlation values indicate better representativeness. In the hierarchical clustering analysis, we utilized significantly differentially expressed genes between poor responders and good responders to perform clustering. In the Spearman correlation analysis, we approached from another perspective, using the 2,000 most varied genes in cell lines to identify the similarity between cell lines and poor responders. Whether conducting similarity assays with baseline samples or post-chemotherapy samples, among the 21 TNBC cell lines, 10 cell lines consistently ranked within the top 100 in correlation values. We believe that these cell lines can better represent the gene profiles of TNBC poor responders both before and after treatment.

The single-sample Gene Set Enrichment Analysis (ssGSEA) is a powerful computational method to evaluate the activity of biological pathways and gene sets within individual samples. It quantifies the relative enrichment of predefined gene sets or pathways by considering the expression levels of genes within each sample. In our study, we selected 50 Molecular Signatures Database (MSigDB) hallmark gene sets [20] for ssGSEA calculations. Many of these hallmark gene sets directly relate to cancer development and progression, including cell cycle regulation, apoptosis, DNA repair, epithelial-mesenchymal transition (EMT), and hypoxia. Assessing these gene sets allows us to analyze better which cell lines exhibit more significant similarity with TNBC chemo poor responders. This analysis offers valuable insights into the functional similarities between cell lines and patient samples, helping to identify more accurate and relevant models for studying TNBC poor responders based on their pathway activation patterns (Figures 1D and 2D).

In summary, to comprehensively and accurately identify our study’s most representative cell lines, we employed Hierarchical clustering analysis, Spearman’s rank correlation, and GSEA analysis to determine the representative cell lines based on gene expression levels and pathway activation patterns. In the similarity analysis with baseline TNBC poor responder samples, five cell lines were found to be the most representative across all three methods. These include MDA-MB-231, MDA-MB-157, HCC1187, HCC38, and HCC1395. In the similarity analysis with post-chemo TNBC poor responder samples, three cell lines, namely MDA-MB-231, MDA-MB-157, and HCC1395, were consistently significant across all three methods.

Notably, MDA-MB-231, MDA-MB-157, and HCC1395 not only serve as the most representative cell lines for TNBC poor responders at the baseline time point but also represent the most suitable models for post-chemotherapy TNBC poor responders. These cell lines could provide valuable insights into the molecular mechanisms driving TNBC poor response to chemotherapy and help facilitate the development of more effective treatment strategies. Therefore, in our following experiment, MDA-MB-231 cell line were selected for following research.

## Materials and Methods

### Data Collection and Integration

In our previous study, we searched the Gene Expression Omnibus (GEO) [21] using the keywords “breast cancer” and “chemo” to conduct pathway and sub-pathway analyses, revealing the activation of several molecular pathways associated with chemoresistance in breast cancer patients [21]. After manual review and considering the Miller-Payne (MP) score, a grading system reflecting patients’ drug resistance status, we selected relevant data and developed cohorts of good and poor chemotherapy responders. TNBC is a subtype of breast cancer, and in our current studies, we have chosen TNBC data from our previous data collections. We also obtained genomic profiles of 21 TNBC cell lines from The Cancer Cell Line Encyclopedia (CCLE) platform. In CCLE database, it has characterized the genomic and transcriptomic profiles of over 900 cell lines [22]. Dataset descriptions and sample sizes for each dataset are displayed in Table 1. All samples were generated using the same Affymetrix Human Genome U133 Plus 2.0 Array platform.

**Table 1.** Genomic dataset in breast cancer chemo-resistant study and genomic profile of CCLE cell line.

| Dataset | Sample sizes | Platform | Source | Publication | Genes and Pathways Discovered in Breast Cancer Chemo Resistant |
| --- | --- | --- | --- | --- | --- |
| GSE28844 | 61 | Affymetrix | Patient | Laura et al. 2013 [32] | chemo resistant associated gene shows enriched in Wnt, HIF1, p53, and Rho GTPases signaling pathways. |
| GSE18728 | 61 | Affymetrix | Patient | Korde et al, 2010[33] | MAP-2, MACF1, VEGF-B,EGFR showed up-regulated in poor responder after chemotherapy |
| GSE32646 | 115 | Affymetrix | Patient | Miyake T et al,2012[34] | GSTP1 expression predicts poor response of neoadjuvant chemotherapy in ER-negative breast cancer patient |
| GSE36133 (CCLE) | 917 (21 TNBC) | Affymetrix | Cell line | Barretina et al, 2010[22] | . |

### Cell line similarity analysis

#### Data pre-processing for gene expression profiles

For microarray data acquired from the GEO database, we conducted RMA normalization using Expression Console™ (EC) software version 1.1 (Affymetrix). The signal intensity was logged with base 2 to stabilize the variance. Next, we filtered the microarray data based on the following criterion: if 80% of the samples had expression levels for each individual lower than the background level. Affymetrix probe IDs were mapped to gene symbols based on their GPL platforms. When multiple probes mapped to a single gene, the median expression value was used in the analysis.

#### Different gene expression and Hierarchical Clustering analysis

The Limma package [23] [24]in R software (version 3.6.1) was employed for differential gene expression (DGE) analyses. We adjusted nominal P-values using the Benjamini-Hochberg method and calculated fold change by comparing mean expressions between non-response and response groups. Hierarchical clustering is a widely used algorithm in bioinformatics that groups similar objects [25, 26] performed a hierarchical clustering analysis between CCLE TNBC cell lines and TNBC patients using significantly upregulated and downregulated genes.

#### Exploring correlation between TNBC cell lines and TNBC poor responder samples

In the CCLE dataset, there are gene expression values for over 10,000 genes across 916 cell lines. Firstly, we calculate the standard deviation of each gene across the 916 cell lines to understand the variation of these genes in all CCLE cell models Most 2000 variation genes in this cell lines were select for the Spearman correlation analysis. For each cell line, average Spearman correlation coefficient across all samples of TNBC poor responder. Then the Higher average correlation values indicate better representativeness. The CCLE dataset contains gene expression values for over 10,000 genes across 916 cell lines. To assess the representativeness of these cell lines, we first calculate the standard deviation for each gene across the 916 cell lines, which allows us to evaluate the variation of these genes among all CCLE cell models. Subsequently, we select the top 2,000 most variable genes in these cell lines to perform the Spearman correlation analysis. Finally, for each cell line, we calculate the average Spearman correlation coefficient across all samples of TNBC poor responders, which serves as a measure of similarity between the cell line and the patient samples. Higher average correlation values indicate better representativeness of the cell line model for the TNBC poor responder population.

#### ssGSEA pathway similarity analysis

ssGSEA scores were obtained by R package “GSVA”[27]for or the 50 MSigDB hallmark gene sets [28]. Hierarchical clustering analysis was performed to explore the similarity between cell lines and TNBC poor responder samples in the context of biological processes and pathways, utilizing the ssGSEA score.

#### GO and KEEGG pathway analysis

Gene Ontology (GO) [29]and Kyoto Encyclopedia of Genes and Genomes (KEGG) [30] gene enrichment analyses are widely used approaches for interpreting high-throughput gene expression data. These analyses help identify biological processes, molecular functions, and cellular components (GO) or metabolic and signaling pathways (KEGG) significantly enriched for a given gene set. Our study utilized the R package “clusterProfiler” [31] to perform GO and KEGG pathway enrichment analyses. We set the threshold at an adjusted P-value of less than 0.05 and selected the top 20 pathways for further investigation.

